# Positive correlations in susceptibility to a diverse panel of viruses across host species

**DOI:** 10.1101/2024.10.09.617458

**Authors:** Ryan M. Imrie, Megan A. Wallace, Ben Longdon

**Affiliations:** Centre for Ecology & Conservation, Faculty of Environment, Science, and Economy, University of Exeter, Penryn Campus, Penryn, United Kingdom; MRC-University of Glasgow Centre for Virus Research, Glasgow G61 1QH, UK

## Abstract

Our ability to predict the emergence of novel viruses relies on there being generalisable patterns in the susceptibilities of hosts to novel infections. Studies investigating variation in susceptibility among host species have consistently shown that closely related hosts share similar susceptibilities to a given virus. However, the extent to which such phylogenetic patterns of susceptibility are correlated amongst diverse sets of viruses is unclear. Here, we investigate phylogenetic correlations in susceptibility among *Drosophilidae* hosts to a panel of eleven different invertebrate viruses, comprising seven unique virus species, six unique families, and both RNA and DNA viruses. The susceptibility of hosts to each pair of viruses tested was either positively correlated across host species or did not show evidence of correlation. No negative correlations, indicative of evolutionary trade-offs in host susceptibility to different viruses, were detected between any virus pairs. The strength of correlations were generally higher in viruses of the same species and family, consistent with virus phylogenetic patterns in host infectivity. Our results suggest that generalised host susceptibility can result in positive correlations, even between highly diverged viruses, while specialised interactions with individual viruses cause a stepwise decrease in correlation strength between viruses from the within-species, to the within-family, to the across-family level.

## Introduction

The emergence of viruses in novel host species continues to pose a significant challenges to public health, with epidemics in humans and animals often occurring as a direct result of virus host shifts, where a virus jumps from one host species to another [1–3]. Despite considerable interest in the growing field of zoonotic risk prediction [4], virus host shifts remain largely unpredictable and difficult to control, as illustrated by the recent and unexpected shift of avian H5N1 into domestic cattle, which has been accompanied by a novel tissue tropism and route of transmission for the virus [5]. It remains an open question to what extent the traits of novel infections can be meaningfully predicted given the diversity of hosts and viruses that exist in nature [6]. However, generalisable patterns in host-virus interactions can provide rules of thumb to help develop frameworks for understanding pathogen emergence.

One such pattern that appears consistently in both experimental and natural studies of infection is the influence of host evolutionary relatedness [7], which can explain a large proportion of variation in virulence [8–10], transmissibility [11], and viral load [12–14] across host species. Host phylogenetic effects, which act as a proxy for divergence in traits (for example physiology and immunity), can result in two distinct patterns. Firstly, “distance effects”, where viruses become progressively less well adapted to a novel host as evolutionary distance between donor and recipient species increases, can result in decreased replication [12], increased virulence [11] and decreased onward transmission [10,11,15]. Secondly, independent of distance to the donor host, groups of closely related hosts present similar environments to novel viruses and thus tend to share similar infection phenotypes which can be referred to as “clade effects” [7].

Host phylogenetic patterns in susceptibility have been described for many viruses, however it is unclear how patterns of host susceptibility to one virus will be correlated with that of another virus [13]. Positive phylogenetic correlations in host susceptibility may occur due to overlap in the genetic components underlying susceptibility to each virus, or if these susceptibilities have a shared evolutionary history [16,17]. Yet, there is good reason to expect that hosts will not have similar levels of susceptibility to all viruses. For example, viruses that use conserved host receptors may be able to enter and infect a diverse range of hosts [18], but the cellular machinery viruses hijack to replicate will vary across host species and amongst viruses, as will the immune responses viruses must evade or suppress, leading to differences in susceptibility. Immune responses may be general, or more specific to certain pathogens or pathogen types. This variation may in part be due to selection by pathogens, which in some instances may lead to reciprocal coevolutionary changes in the pathogens commonly infecting a given host [19]. Non-immunity mediated resistance (*e*.*g*., alterations to pro-viral factors) may be specific to certain pathogen genotypes and species or may have effects on a broader range of pathogens [20]. On the other hand, pathogens may fail to infect a given host due to their specialisation on another host, having adapted to diverged components of their “natural” host’s physiology, cellular machinery, and immune responses [19].

While some similarities have been observed between viruses at broad taxonomic scales – such as viruses belonging to the same family having similar propensities for cross-species transmission [15,21,22] – few studies have formally tested for correlations between viruses in their ability to infect across host species. This is likely due to both the scale of the experiments required to measure these correlations, and the often limited number and diversity of virus isolates available for study. As such, it remains unclear how common correlations in these patterns are between viruses, or what factors may produce or influence them. In a previous study, we have shown that two invertebrate *Dicistroviruses*, Drosophila C virus (DCV) and Cricket Paralysis virus (CrPV) are positively correlated in their phylogenetic patterns of susceptibility across *Drosophilidae* host species. The strength of correlation between these two virus species (r ≈ 0.55) was weaker than the correlations that exist between different natural strains of DCV (r ≈ 0.95), suggesting that the extent to which *Dicistroviruses* share the same host phylogenetic pattern of susceptibility decreases as they become more evolutionarily distant [13]. Here, we expand on these findings by estimating the phylogenetic correlations that exist in virus susceptibilities across a larger and more diverse insect virus panel, allowing for comparisons within virus species, within virus family, across virus family, and across nucleic acid type. A total of eleven viruses are included in this study, encompassing seven unique species, six families, and including both RNA and DNA viruses (Table 1).

**Table 1:**
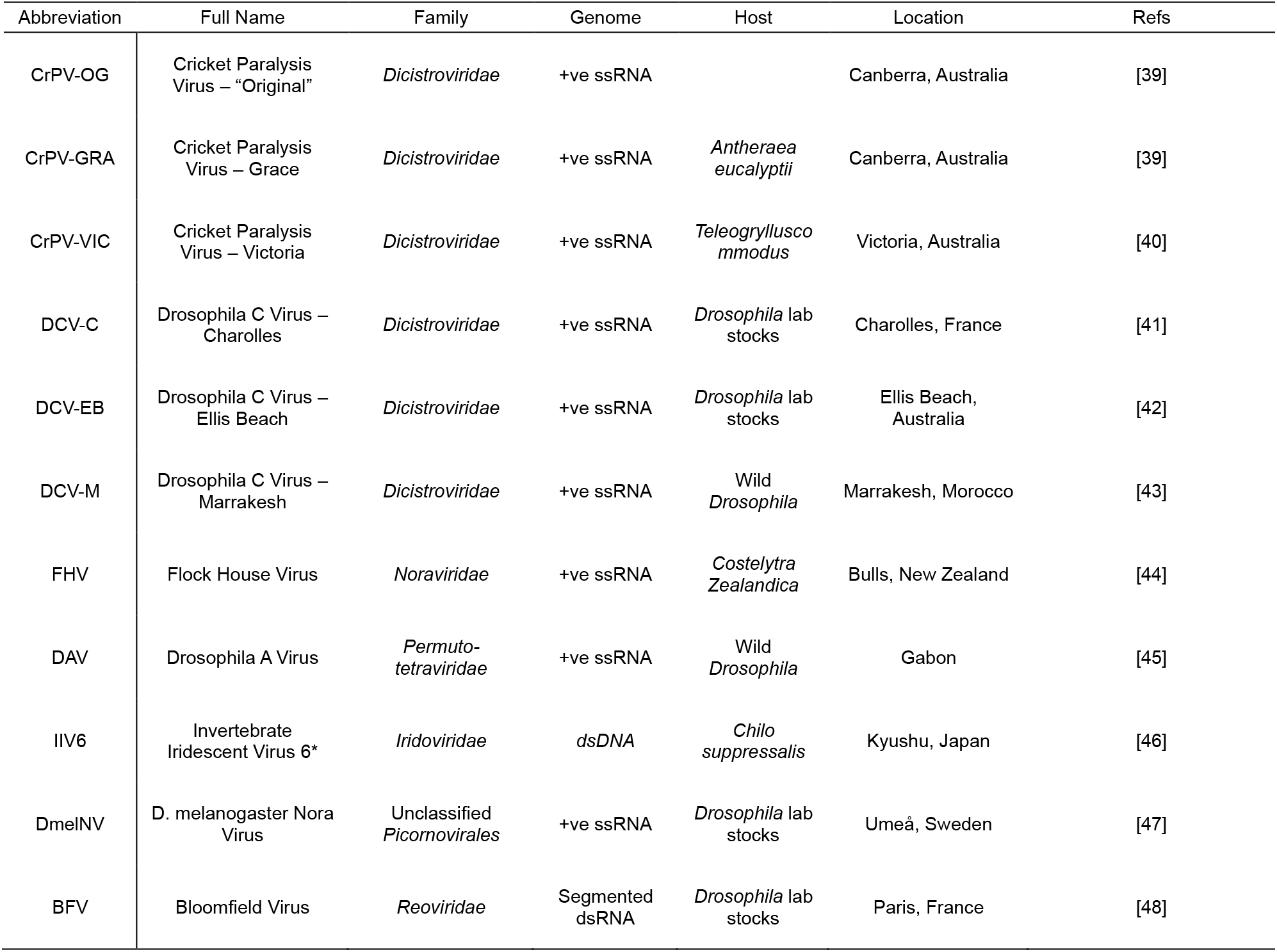
Identities of virus isolates used in this study. Cricket Paralysis Virus – “Original” is the typical CrPV isolate used in most studies of this virus species to date. Phylogenetic reconstruction of the available CrPV isolates suggests CrPV-OG most likely originated as an aliquot of CrPV-BEE (supplementary Figure 1) and has since undergone an unknown number of passages in laboratory host stocks. (*) Also referred to as Chilo Iridescent Virus [49].

Variation in immune responses among host species may provide a basis for phylogenetic correlations between viruses. The majority of our knowledge of antiviral immunity in *Drosophilidae* comes from studies of *Drosophila melanogaster* which have described a small number of generalised immune pathways, namely the antiviral RNAi pathway; the antimicrobial signalling pathways Toll, IMD, JAK-STAT, and STING; and the processes of virally stimulated autophagy and phagocytosis [23–26]. Viral suppression of these generalised host immune pathways is common and can vary across hosts (Supplementary Table 1)[27–32]. For example, the activity of Drosophila Nora Viruses (NVs) antiviral RNAi suppressors is host species specific [30]. Variation in the ability of different viruses to effectively supress host immunity across host species could potentially mask general susceptibility to certain groups of viruses. In addition, several genes have been described that confer specific immunity to some of the viruses included in this study: variation in the *Pastrel* and *Ubc-E2H* genes have a large effect on DCV and CrPV susceptibility but not Flock House Virus (FHV) [33,34], and a TE-mediated truncation of the *Veneno* gene influences Drosophila A Virus (DAV) susceptibility but not DCV or FHV [35].

Since *Drosophila* antiviral immunity comprises both generalised and specialized components, we may expect positive phylogenetic correlations to exist between viruses due to overlap in the host genetic components influencing susceptibility, with correlations strengths shifting towards zero for viruses with specialised host interactions. Previous studies that have compared host susceptibilities to multiple *Drosophila* viruses found positive correlations in susceptibility across both genotypes of *D. melanogaster* and different host species, although viruses tested are typically closely related [12,13,33,36]. These studies also provide evidence of pairs of viruses that show no detectable correlations. Interestingly, no study has reported a significant negative correlation between viruses, which would indicate from a trade-off where increased resistance to one virus came at the cost of decreased resistance to another virus. The existence of trade-offs and positive correlations have broad implications for the evolution of immunity, as any non-independence in susceptibility can influence the genetic diversity, the strength and direction of selection, and community compositions of both the host and viruses [16,37,38].

## Materials & Methods

### Fly Species

Laboratory stocks of 35 species of *Drosophilidae* (Supplementary Table 2) were maintained in multi-generation stock bottles at 22°C with a 12-hour light-dark cycle. Each bottle contained 50ml of one of four varieties of media, which were chosen to optimise rearing conditions (for media recipes see http://doi.10.6084/m9.figshare.21590724.v1). While these food varieties vary in macronutrient content, adult diet has been shown to have little effect on the outcome of viral infection in this system [50]. Inference of the host phylogeny has been described in detail elsewhere [8]. Briefly, publicly available sequences of the *28S, Adh*,

*Amyrel, COI, COII, RpL32*, and *SOD* genes were collected from Genbank (see https://doi.org/10.6084/m9.figshare.13079366.v1 for a full breakdown of genes and accessions by species). Gene sequences were aligned in Geneious v9.1.8 (https://www.geneious.com) using a progressive pairwise global alignment algorithm with free end gaps and a 70% similarity IUB cost matrix. Gap open penalties, gap extension penalties, and refinement iterations were kept as default.

Phylogenetic reconstruction was performed using BEAST v1.10.4 [51] as the subsequent phylogenetic mixed model (see below) requires a tree with the same root-tip distances for all taxa. Genes were partitioned into separate ribosomal (*28S*), mitochondrial (*COI, COII*), and nuclear (*Adh, Amyrel, RpL32, SOD*) groups. The mitochondrial and nuclear groups were further partitioned into groups for codon position 1+2 and codon position 3, with unlinked substitution rates and base frequencies across codon positions. Each group was fitted to separate relaxed uncorrelated lognormal molecular clock models using random starting trees and 4-category gamma-distributed HKY substitution models. The BEAST analysis was run twice, with 1 billion MCMC generations sampled every 100,000 iterations, using a birth-death process tree-shape prior. Model trace files were evaluated for chain convergence, sampling, and autocorrelation using Tracer v1.7.1 [52]. A maximum clade credibility tree was inferred from the posterior sample with a 10% burn-in.

### Virus Isolates

The eleven virus isolates used in this study were kindly provided by Julien Martinez (DCV isolates) [53]; Karyn Johnson (CrPV-GRA and CrPV-VIC) [54]; Jon Day and Frank Jiggins (DAV, FHV) [35]; and Jared Nigg, Valérie Dorey, and Maria Carla Saleh (CrPV-OG, DmelNV, BFV, IIV6) [55]. Prior to infection, the concentrations of viral RNA in each stock were measured using qRT-PCR (described below) and normalised by dilution with Ringers solution [56]. Each stock was also checked via qRT-PCR for contamination with each of the other viruses included in this study, with no contamination detected.

Multiple DCV and CrPV isolates are available for experimental study [53,54], and the isolates included here were chosen to represent distinct sub-clades within the *Dicistrovirus* phylogeny (Supplementary Figure 1). To infer this phylogeny, genome sequences of the DCV isolates were downloaded from Genbank (see Supplementary Table 3 for accessions), and CrPV genomes were sequenced as follows. Separate vials of 20 7-8 day old male *D. melanogaster* flies were experimentally inoculated with each CrPV isolate (see Inoculation below), and the viruses allowed to amplify for 2 days before being snap frozen in liquid nitrogen, homogenized in Trizol (Invitrogen), and RNA extracted using phenol-chloroform phase separation. RNA quantity was assessed using the Qubit RNA BR kit (ThermoFisher) and checked for integrity using the Agilent RNA 6000 Nano kit and the Agilent 2100 Bioanalyzer. Strand-specific 150bp paired-end total RNA libraries were prepared for each sample with Ribo-zero Plus (Illumina) rRNA depletion, and RNAseq was performed on an Illumina NovaSeq 6000 SP flow cell to a depth of >20M reads per sample.

Raw reads were processed using Cutadapt [57] to remove adaptors, FastQC [58] to filter by overall quality and length, and Trimmomatic [59] to quality trim the ends of reads, all with default settings. Processed reads were then *de novo* assembled into contigs using Trinity [60]. To identify contigs belonging to *Drosophila* viruses, the longest open reading frame of each contig was translated and queried against a *Drosophila* virus database maintained by

Darren Obbard (https://obbard.bio.ed.ac.uk/data.html) using BLASTn [61] and DIAMOND [62]. BLAST hits with >60% similarity at the nucleotide level were retained to allow for the identification of any diverged variants. Contigs matching known *Drosophila* viruses were manually inspected for contaminants and fly genomic material. All virus sequences can be found on Genbank with the accession numbers PQ246907 to PQ246914 (Supplementary Table 3).

To infer the *Dicistrovirus* phylogeny, coding and non-coding sequences of CrPV and DCV were separated and aligned in MUSCLE 5.1 [63] using default settings, with translation-alignment used for coding sequences. Two concatenated alignments – one of coding sequences and the other of non-coding sequences – were then used as separate partitions in a phylogenetic reconstruction using BEAST version 1.10.5 [51]. Each partition was fitted to a separate uncorrelated relaxed lognormal molecular clock model with a speciation birth-death process tree-shaped prior [64]. Separate HKY substitution models were used for each partition with a four-category gamma distribution rate of variation, and the coding sequences were further partitioned into separate groups for codon positions (1+2) and (3). Models began with random starting trees, and were run for 2 billion MCMC generations, sampled every 200,000 iterations. The model output was evaluated for convergence using Tracer version 1.7.1 [52], and a maximum clade credibility tree was inferred from the posterior sample with a 10% burn-in.

### Inoculation

Prior to inoculation, 0-1 day old male flies were collected and housed in vials with cornmeal media at 22°C and 70% relative humidity, under a 12-hour light-dark cycle. Each vial contained between 7 and 15 flies (mean = 13.6, Supplementary Table 2). Flies were transferred to fresh media every 2 days until reaching 7-8 days old, at which point they were anaesthetised with CO_2_ and experimentally infected with virus inoculum via pin prick with a 12.5μm diameter stainless steel needle (Fine Science Tools, CA, USA). Each needle was bent to a right angle approximately 250μm from the tip to create a depth stop, and the needle inserted up to this point into the right lateral anepisternal cleft of each fly to provide consistent inoculations. This inoculation route bypasses the gut’s immune barrier but avoids differences in dosage that can occur due to different feeding behaviours across the host species [65]. Male flies were used exclusively to avoid any confounding effects of sex or mating status, which can introduce additional variation in susceptibility to pathogens in female flies [66–68], and recent experiments have shown that male and female flies are strongly positively correlated in their viral susceptibilities across host species [69].

### Measuring Change in Viral Load

At 2 days (±2 hours) post-inoculation, flies were snap frozen in liquid nitrogen and homogenized in Trizol using 0.2mm Zirconia beads (Thistle Scientific) on an Omni Bead Ruptor 24 (4m/s for 15 seconds). Total RNA was extracted using a chloroform-isopropanol extraction method and reverse transcribed to cDNA using Promega GoScript reverse transcriptase with random hexamer primers (Sigma). For IIV6 infected flies, DNA was extracted by removing the RNA-containing upper aqueous phase of the phenol-chloroform extraction and adding a back-extraction buffer of 4M guanidine thiocyanate, 50mM trisodium citrate, and 1M TRIS. The new DNA-containing upper aqueous phase was then collected, and DNA precipitated with isopropanol overnight at -20°C. The resulting pellet was washed three times with 70% ethanol, resuspended in 8mM NaOH, and pH neutralised with 1M HEPES.

q(RT)-PCR was performed for each virus and the housekeeping gene *RPL32* using the Sensifast Lo-Rox SYBR kit (Bioline) on an Applied Biosystems QuantStudio 3 (for primers and cycling conditions, see Supplementary Tables 4-7). Two technical replicates were performed for each reaction and sample, with between-plate variation in C_t_ values corrected for statistically using previously described methods [70,71]. Amplification of the correct products was verified by melt curve analysis, with ±1.5°C and ±3°C used as inclusion cut-offs for viral and RPL32 amplicons respectively, and technical replicates averaged to produce mean virus and RPL32 C_t_s.

Inoculation doses for each virus were collected in *D. melanogaster* by snap freezing eight vials of 15 males per virus immediately after inoculation and processing as above. These values were then used to infer the RPL32-normalised inoculation as:

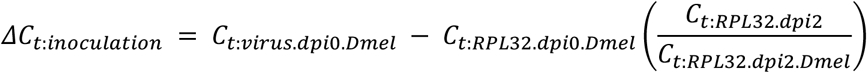

Applying this method to data from a previous study where inoculation doses were collected for each individual species showed the above method was able to accurately infer normalised inoculation doses for each species (R^2^ = 0.92). Fold-changes in viral load were then calculated for each biological replicate using the 2^-ΔΔCt^ method, where *ΔΔC*_*t*_ = *ΔC*_*t:inoculation*_ − *ΔC*_*t:dpi2*_.

### Phylogenetic Modelling and Interspecific Correlations

Phylogenetic generalised linear mixed models were fitted to log_10_-transformed fold-changes in viral load using the R package MCMCglmm [72]. To provide estimates of phylogenetic heritability, univariate models were fitted for each virus, with random effects of both the host phylogeny and a species-specific random effect that explicitly estimated the non-phylogenetic component of variation in viral loads. The proportion of among species variance that can be explained by the host phylogeny – equivalent to phylogenetic heritability, or Pagel’s lambda – was calculated as 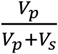, where *V*_*p*_ and *V*_*s*_ are the phylogenetic and species-specific components of variance in viral loads [73–75]. Occasionally, these models struggled to separate the phylogenetic and non-phylogenetic components of variation, and so additional models without the species-specific random effect were fitted. These models then provided an estimate of the repeatability of viral load within-species after accounting for any phylogenetic variance, calculated as 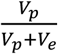, where *V*_e_ is the residual variance of the model [76].

To measure phylogenetic correlations between pairs of viruses, bivariate versions of the above models were fitted with a fixed effect of each virus isolate and 2x2 covariance structures on the random effects and residuals. The phylogenetic covariance matrices were then used to calculate the coefficients of the phylogenetic correlations between viruses as 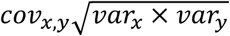, and the regression slopes as 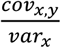. The intercepts of each regression were calculated from the fixed effect estimates of the across-species means and the regression slopes as 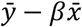. Correlations between viruses were considered significant if the 95% credible intervals (CIs, calculated as the 95% highest posterior density intervals throughout this study) for both the correlation coefficient and slope of the relationship did not overlap with zero. Point estimates for all effects are presented as posterior means.

Models were run for 13,000,000 iterations, with a 3,000,000 iteration burn-in, and were sampled every 5,000 iterations. The results presented here are from models with parameter-expanded priors placed on the covariance matrices, and inverse-gamma priors placed on the residual variances. To ensure our results were robust to changes in prior distribution, models were also fitted with flat and inverse-Wishart priors, which provided qualitatively similar results. As all fly and molecular work was completed by a single researcher, no correction for technical variation added by individual was included.

Full details on the structure of each of the above models can be found in Supplementary Methods, and all scripts and data used in this analysis are available at https://github.com/ryanmimrie/Publications-2024-Drosophilidae-Virus-Phylogenetic-Correlations.

## Results

To determine the strength of phylogenetic correlations in susceptibility to different viruses across *Drosophilidae* we experimentally infected 35 host species with eleven different virus isolates, comprising seven unique virus species, six virus families, and including nine +ssRNA viruses, a dsRNA virus, and a dsDNA virus. Susceptibility, measured as the fold-change in viral load after 2 days of infection, showed considerable variation across host species and virus isolates (Figure 1). Apart from BFV, which persisted at detectable levels in a few host species but failed to amplify to a higher viral load than the inoculation dose in all cases, all viruses showed an ability to persist and replicate in the majority of host species tested. Almost all viruses showed strong evidence of phylogenetic patterns in susceptibility (Figure 2) with the majority of across host species variance in viral load explained by the host phylogeny (Supplementary Table 8).

**Figure 1:**
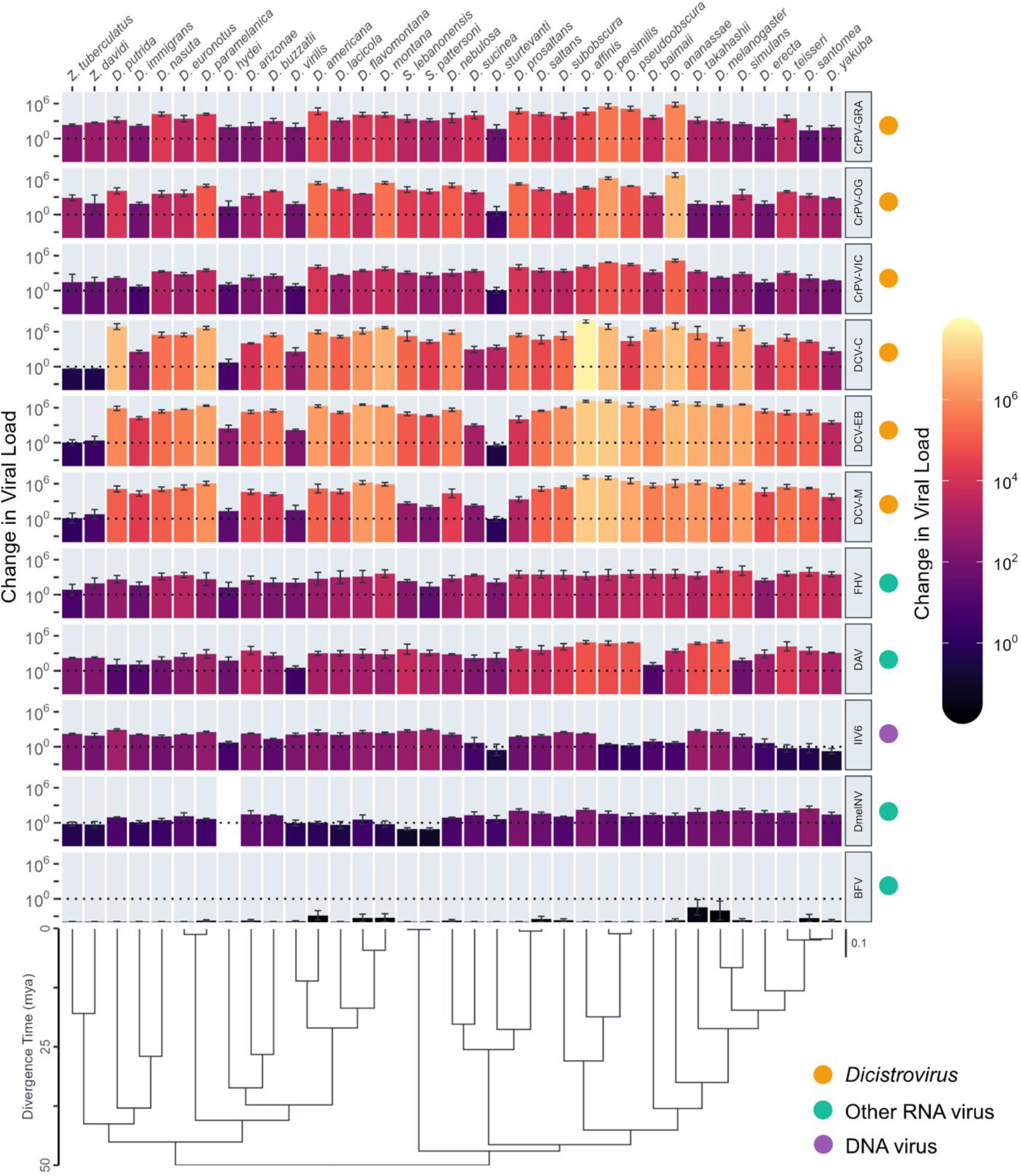
Fold-changes in viral load across *Drosophilidae* host species after infection with different viruses. Bar heights and colour show the mean change in viral load from 0 to 2 dpi on a log_10_ scale, with error bars representing the standard error of the mean. The phylogeny of *Drosophilidae* hosts is shown on the left, with a scale bar for the number of nucleotide substitutions per site and an axis showing the approximate age since divergence (mya) taken from estimates in [77]. Due to off-target amplification, no viral load data was collected from *D. hydei* for DmelNv infection.

**Figure 2:**
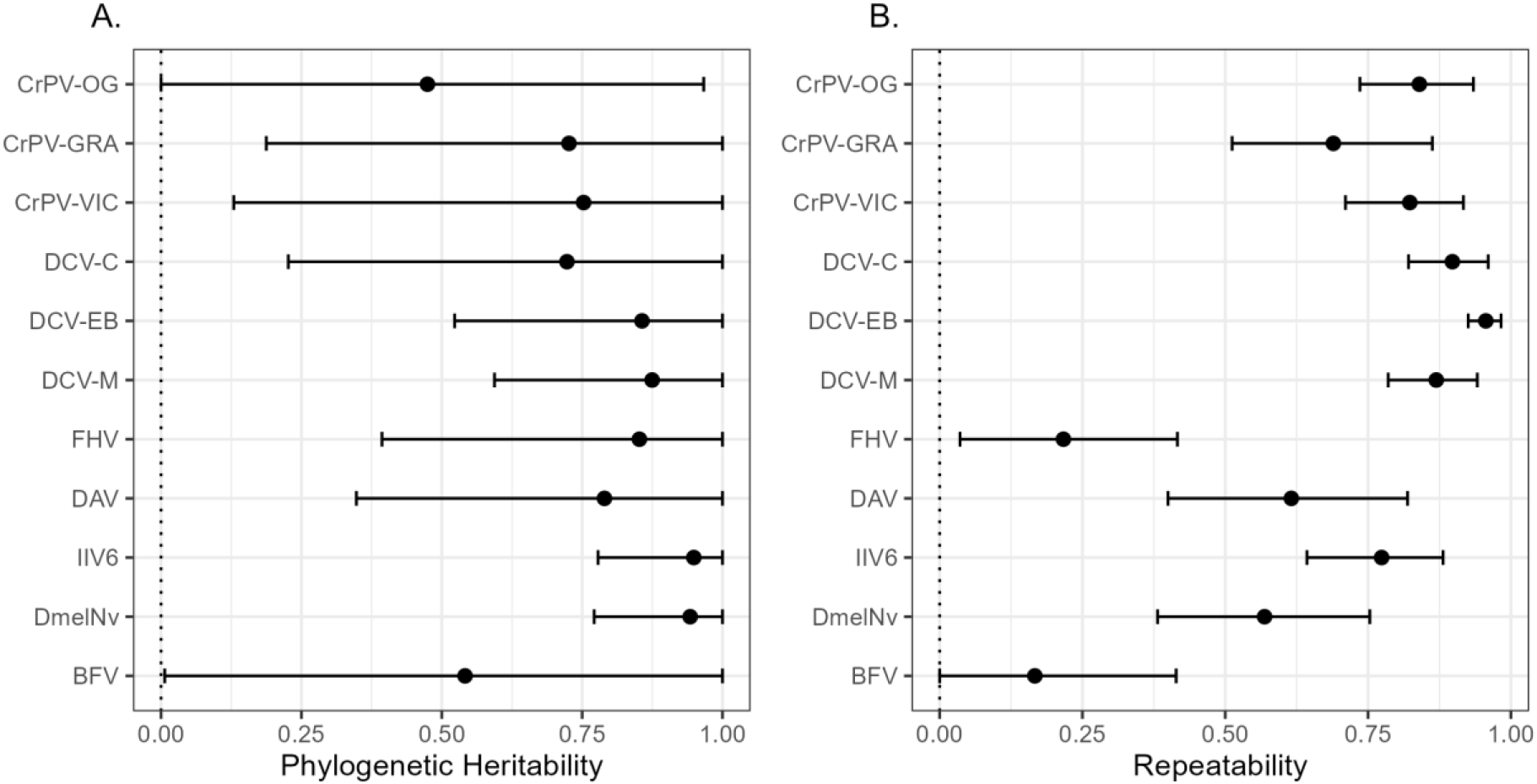
Estimates of phylogenetic heritability and repeatability in viral load for each virus. Values for the phylogenetic heritability (A) and repeatability (B) of fold-changes in viral load were taken from univariate models with (A) or without (B) the inclusion of a non-phylogenetic species-specific random effect. Points represent the mean, and error bars the 95% of the posterior distributions of each estimate (see Table S8 for values).

When we examined the phylogenetic correlations that exist between different viruses, we found that almost all (54/55) point estimates of the correlation coefficients were positive, with 30/55 being significantly positive, and the remaining correlations all indistinguishable from zero (Figure 3). Across our data, we detected no negative phylogenetic correlations in virus susceptibility. Strong positive phylogenetic correlations were detected between all CrPV and DCV isolates, consistent with previous studies of these viruses [13,36], and correlation coefficients were generally higher and more tightly estimated within each virus species than between them. However, with a few exceptions, the 95% credible intervals of the within and between species correlation coefficients overlapped, and so evidence here for there being stronger phylogenetic correlations within virus species is inconsistent. Outside of the *Dicistrovirus* clade, 15/40 virus combinations were positively correlated, 14 of which were between a *Dicistrovirus* and a non-*Dicistrovirus*, including three detectably positive correlations between DCV isolates and the DNA virus IIV6. Only one positive correlation, between FHV and DmelNv, was detected between a non-*Dicistrovirus* pair.

**Figure 3:**
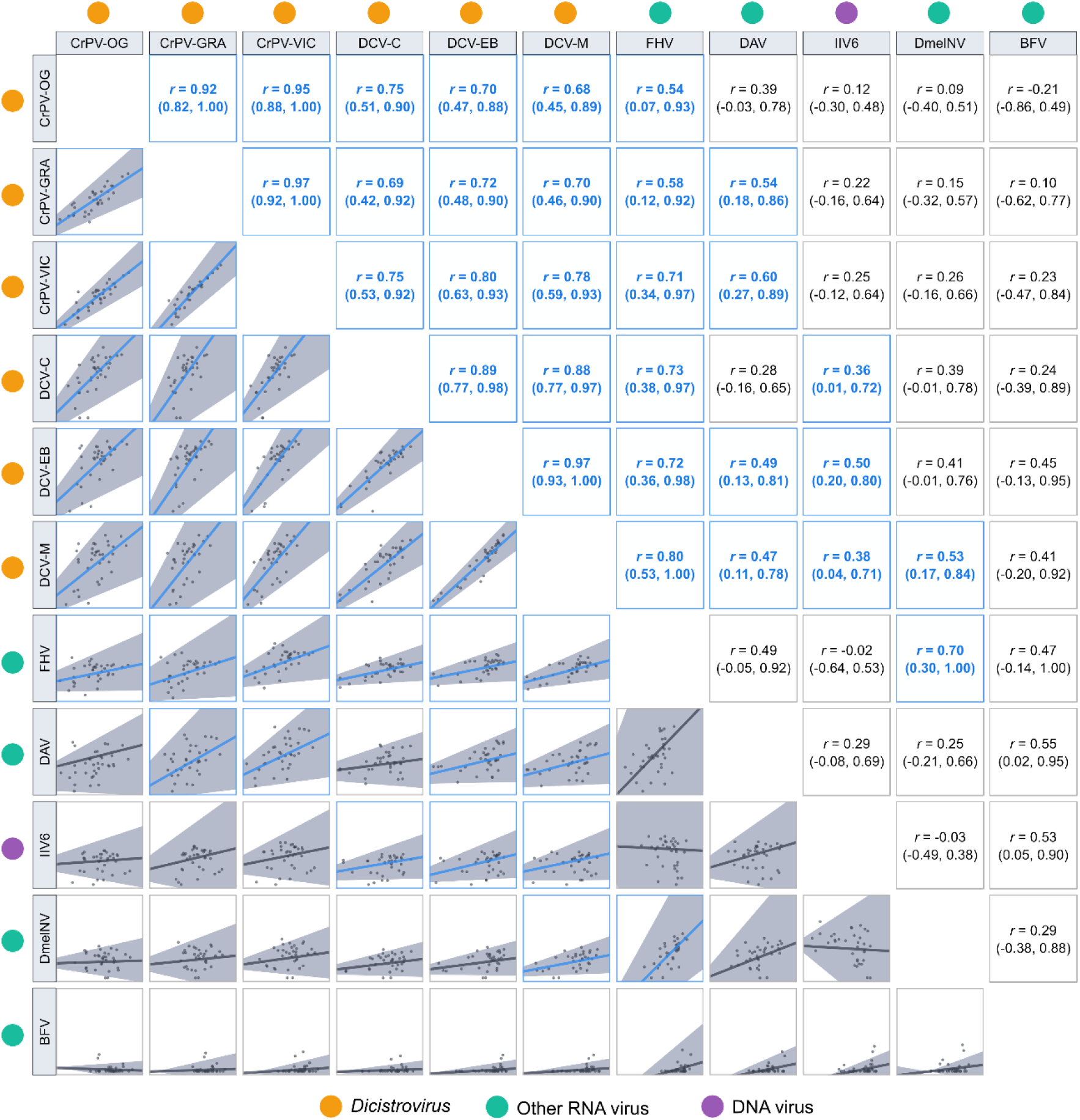
Correlation matrix of fold-changes in viral load across *Drosophilidae* host species infected with different viruses. Points represent the mean viral load at 2dpi for each host species on a log_10_ scale. Interspecific correlation coefficients (r) and trend lines are taken from bivariate versions of model (2), with 95% credible intervals shown in brackets and grey shaded areas. Correlations with coefficient estimates significantly different from zero are highlighted with blue trend lines.

When correlation estimates were grouped depending on if the pair of viruses was from the same species, same family, or different families, a stepwise pattern emerged where the within-species correlations were significantly stronger (r = 0.93, 95% CI: 0.82, 0.99) than the across-family correlations (r = 0.33, 95% CI: -0.18, 0.79) and the within-family correlation estimates (r = 0.73, 95% CI: 0.49, 0.93) overlapped with both other groups (Figure 4). This finding was robust to the inclusion or exclusion of BFV from this analysis (Supplementary Tables 9-10).

**Figure 4:**
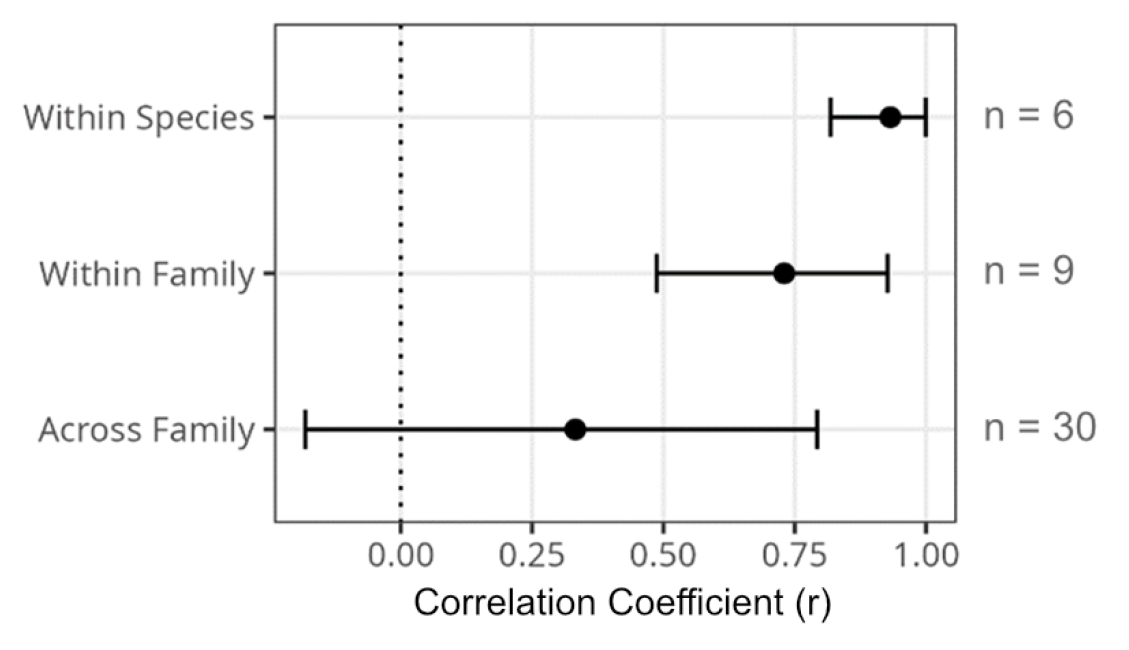
Combined estimates of the strength of correlations between viruses at different evolutionary scales. Posterior distributions of the correlation coefficient estimates for each virus pair were combined to create overall distributions for estimates within virus species, within virus family, and across virus family. Points represent the means, and error bars the 95% CIs of these combined posterior distributions (see Table S9-10 for values).

## Discussion

In this study, we have found that the majority of host phylogenetic correlations in susceptibility to different viruses are positive. Noticeably, no pairs of viruses were negatively correlated in their patterns of susceptibility across host species, suggesting that resistance to one virus generally increases resistance to other viruses. Every virus provided evidence of at least one positive phylogenetic correlation with another virus except BFV, which was undetectable at 2 dpi in most of the host species tested here.

Three general categories of correlation appear to exist between the viruses tested here: strong positive phylogenetic correlations, which are apparent between all *Dicistrovirus* pairs, and between each DCV isolate and FHV; weaker positive phylogenetic correlations, such as those between the *Dicistroviruses* and DAV, and between DCV and IIV6; and viruses with no detectable phylogenetic correlation, such as CrPV and IIV6, and DmelNV and DAV. The strong positive correlations within the *Dicistroviruses* may be explained by the relatively high degree of similarity between DCV and CrPV genomes (57-59% nucleotide and amino acid similarity [13]), which likely causes them to share many of the same host interactions during infection. Evolutionary relationships between viruses do not explain the patterns in our data when compare more distantly related viruses outside of the *Dicistroviruses*. DCV, CrPV, and DmelNv are all members of the order *Picornavirales*, and yet the *Dicistroviruses* show more consistent evidence of positive correlations with FHV and DAV, which converge with *Dicistroviruses* only at the higher taxonomic levels of kingdom (*Orthornavirae*) and realm (*Riboviria*) respectively.

When making comparisons between viruses across larger taxonomic distances such as these, it may be more beneficial to consider the functional homology between viruses instead of their evolutionary similarity. For example, the strong positive correlations between the *Dicistroviruses* and FHV may be better explained by the high degree of overlap in the host immune pathways that control them; RNAi, Toll, IMD, JAK-STAT and phagocytosis have all been implicated in host susceptibility to these viruses, while several of the other viruses included here appear to have escaped the influence of at least one of these host defences.

Functional group/guild modelling of pathogen interactions with host immunity has previously been used as a simplifying approach to successfully infer the outcome of coinfections [78], and it is possible that a similar approach could be used to explain differences in correlation strength between evolutionarily diverged viruses.

The lack of negative phylogenetic correlations could suggest that, as *Drosophila* immune systems have evolved in response to infection, they have not been constrained by a trade-off where increased resistance to one virus has decreased resistance to another virus. Experimental evolution, considered the gold standard of proof for genetic correlations [16], could be used to investigate the ability of virus susceptibilities in *Drosophilidae* to be decoupled, and so offer further insights into the origins of the phylogenetic correlations we have measured. A similar approach has already been used to explore the evolution of virus susceptibility within *D. melanogaster* [34], which showed that selecting for DCV resistance also increased resistance to CrPV and FHV, suggesting these traits do indeed overlap genetically.

Nevertheless, our data suggest that the majority of correlations in virus susceptibilities in *Drosophilidae* are positive. As a model system for across-species infections, our finding suggest that the host phylogenetic patterns of susceptibility to one virus may provide useful information on the patterns of susceptibility of closely related viruses, but this ability to extrapolate between viruses deteriorates in a stepwise fashion with increasing virus taxonomic divergence. Our results imply that accurately predicting the phenotypes of novel viruses for which data on close relatives is not available is likely to be a major challenge that would require a detailed mechanistic understanding of the underlying determinants of virus susceptibility.

## Acknowledgements

We would like to thank the Exeter Sequencing Service for their support, and Julien Martinez, Karyn Johnson, Jon Day, Frank Jiggins, Jared Nigg, Valérie Dorey, and Maria Carla Saleh for providing virus isolates. This project utilised equipment funded by a Wellcome Trust Multi-User Equipment Grant (award number 218247/Z/19/Z). BL, RI, and MW were supported by a Sir Henry Dale Fellowship jointly funded by the Wellcome Trust and the Royal Society (grant number 109356/Z/15/Z) https://wellcome.ac.uk/funding/sir-henry-dale-fellowships. For the purpose of Open Access, the author has applied a CC BY public copyright licence to any Author Accepted Manuscript version arising from this submission.

## Notes

https://github.com/ryanmimrie/Publications-2024-Drosophilidae-Virus-Phylogenetic-Correlations

### Competing Interest Statement

The authors have declared no competing interest.

