## Supplementary Information for "Positive correlations in susceptibility to a diverse panel of viruses across host species"

### Supplementary Materials

The following document contains supplementary methods, tables, and figures for Imrie et al., "Positive correlations in susceptibility to a diverse panel of viruses across host species" (2024).

#### Supplementary Methods

##### Phylogenetic Generalised Linear Mixed Model Structures

Three structures of phylogenetic generalised linear mixed models were used in this study: a univariate structure that included a non-phylogenetic species-specific random effect (1), which was used to estimate phylogenetic heritabilities; a univariate structure with this non-phylogenetic random effect removed, which was used to estimate repeatabilities (2); and a bivariate version of (2) was used to estimate phylogenetic correlations between pairs of viruses (3):

$$y_{hi} = \beta_1 + \mu_{p:h} + \mu_{s:h} + e_{hi} \quad (1)$$

$$y_{hi} = \beta_1 + \mu_{p:h} + e_{hi} \quad (2)$$

$$y_{hiv} = \beta_{1:v} + \mu_{p:hv} + e_{hiv} \quad (3)$$

In these models, $y_{hiv}$ is the change in viral load for virus $v$ in the $i^{th}$ biological replicate of host species $h$. The fixed effect $\beta_1$ represents the intercepts for each virus isolate, the random effect $\mu_p$ represents the effects of the host phylogeny assuming a Brownian motion model of evolution, $\mu_s$ represents the non-phylogenetic species-specific random effect, and $e$ represents the model residuals.

Within each of these models the random effects and residuals were assumed to follow a multivariate normal distribution with a centred mean of zero and a covariance structure of $V_p \otimes A$ for the phylogenetic effects, $V_s \otimes I$ for species-specific effects, and $V_e \otimes I$ residuals, where ⊗ represents the Kronecker product. $A$ represents the host phylogenetic relatedness matrix, $I$ an identity matrix and $V$ represents either a 1x1 or 2x2 covariance matrix describing the between-species variances and covariances of changes in viral load for the different viruses. Specifically, the matrices $V_p$ and $V_s$ describe the phylogenetic and non-phylogenetic between-species variances in viral load for each virus and the covariances between them, while the residual covariance matrix $V_e$ describes within-species variance that includes both true within-species effects and measurement errors. Since each biological replicate was tested with a single virus isolate, the covariances of $V_e$ cannot be estimated and were set to zero.

Models were run for 13 million MCMC generations, sampled every 5,000 iterations with a burn-in of 3 million generations. Parameter expanded priors were placed on the covariance matrices, resulting in multivariate F distributions with marginal variance distributions scaled by 1,000. Inverse-gamma priors were placed on the residual variances, with a shape and scale equal to 0.002. To ensure the model outputs were robust to changes in prior distribution, models were also fitted with flat and inverse-Wishart priors, which gave qualitatively similar results.

### Supplementary Tables

| | Antiviral RNAi | | Toll | | IMD | | JAK-STAT | | STING | | Phagocytosis | |
|---|---|---|---|---|---|---|---|---|---|---|---|---|
| | Controls Virus | Suppressed | Controls Virus | Suppressed | Controls Virus | Suppressed | Controls Virus | Suppressed | Controls Virus | Suppressed | Controls Virus | Suppressed |
| **CrPV** | [1–4] | [5] | [6] | | [4,7] | | [8] | | [9] | | [10] | |
| **DCV** | [1–4,11] | [1] | [6] | | [4,12] | | [8,13] | | [14] | | [15,16] * | |
| **FHV** | [2–4,11,17] | [18] | [6] | | [19] | | [19] | | [4] | | [10] | |
| **DAV** | [20] | | | | [21] | | | | [21] | | | |
| **IIV6** | [4,8,22, 23] | [23] | | [24] | | [24] | [25] | | [4] | | [10] | |
| **DmelNV** | [26] | [27] | [6,26] * | | | | [26] | | | | | |
| **BFV** | | | | | | | | | | | | |

**Supplementary Table 1: Known interactions between each virus species included in this study and *D. melanogaster* antiviral immune pathways.** Dark grey boxes indicate immune pathways that control or become suppressed by the virus, while white boxes indicate pathways that do not control or become suppressed by the virus. Light grey boxes indicate combinations of immune pathway and virus where no/insufficient evidence is available. Results shown have been taken from a combination of *in vitro* and *in vivo* experimental studies, references for which are provided in (Supplementary Table 1). Responses that are known to be context dependent (*e.g.,* dose or infection route dependent) are marked with an asterisk (*), and the response most relevant to the infection context of this study is displayed. References for these effects are given inside each cell.

| | CrPV-OG | | | CrPV-GRA | | | CrPV-VIC | | | DCV-C | | | DCV-EB | | | DCV-M | | | FHV | | | DAV | | | IIV6 | | | DmelNv | | | BFV | | |
|---|---|---|---|---|---|---|---|---|---|---|---|---|---|---|---|---|---|---|---|---|---|---|---|---|---|---|---|---|---|---|---|---|---|
| *D. affinis* | 15 | 15 | 14 | 10 | 13 | 15 | 14 | 13 | 14 | 8 | 15 | 12 | 14 | 14 | 13 | 14 | 15 | 13 | 14 | 14 | 12 | 14 | 14 | 12 | 9 | 15 | 15 | 15 | 14 | 15 | 15 | 11 | 15 |
| *D. americana* | 12 | 15 | 15 | 9 | 15 | 15 | 14 | 11 | 14 | 12 | 10 | 15 | 14 | 15 | 15 | 15 | 15 | 12 | 15 | 13 | 14 | 13 | 13 | 14 | 15 | 15 | 15 | 12 | 14 | 15 | 15 | 14 | 14 |
| *D. ananassae* | 13 | 14 | 15 | 14 | 14 | 14 | 15 | 15 | 15 | 15 | 14 | 15 | 13 | 13 | 13 | 12 | 12 | 14 | 15 | 14 | 13 | 11 | 14 | 15 | 10 | 12 | 14 | 14 | 14 | 12 | 14 | 15 | 11 |
| *D. arizonae* | 15 | 14 | 14 | 10 | 15 | 14 | 12 | 14 | 14 | 14 | 13 | 15 | 15 | 15 | 13 | 13 | 15 | 14 | 14 | 15 | 15 | 10 | 13 | 15 | 13 | 15 | 12 | | 15 | 13 | 15 | 15 | 12 |
| *D. baimaii* | 12 | 14 | 10 | 13 | 15 | 14 | 12 | 11 | 11 | 9 | 14 | 14 | 12 | 12 | 12 | 12 | 11 | 14 | 12 | 15 | 12 | 8 | 14 | 14 | 15 | 15 | 11 | 15 | 10 | 12 | 10 | 10 | 12 |
| *D. buzzatii* | 14 | 13 | 14 | 7 | 15 | 15 | 14 | 14 | 10 | 13 | 13 | 14 | 14 | 15 | 15 | 15 | 12 | 15 | 15 | 15 | 14 | 12 | 15 | 15 | 12 | 14 | 14 | 13 | 15 | 13 | 15 | 15 | 15 |
| *D. erecta* | 13 | 13 | 15 | 13 | 14 | 15 | 14 | 15 | 15 | 13 | 13 | | 13 | 13 | 14 | 14 | 11 | 10 | 14 | 13 | 13 | 13 | 15 | 13 | 15 | 11 | 15 | | 13 | 14 | 13 | 14 | 15 |
| *D. euronotus* | 13 | 15 | 14 | 14 | 15 | 12 | 14 | 13 | 15 | 13 | 15 | 13 | 15 | 13 | 10 | 14 | 13 | 15 | 14 | 11 | 14 | 11 | 14 | 15 | 15 | 15 | 14 | | 14 | 14 | 15 | 14 | 15 |
| *D. flavomontana* | 14 | 13 | 13 | 14 | 14 | 15 | 13 | 15 | 15 | 12 | 13 | 15 | 15 | 14 | 14 | 15 | 15 | 14 | 14 | 15 | 15 | 15 | 15 | 15 | 15 | 14 | 15 | 15 | 15 | 14 | 14 | 15 | 15 |
| *D. hydei* | 15 | 15 | 15 | 14 | 15 | 15 | 13 | 15 | 14 | 15 | 14 | 14 | 15 | 14 | 14 | 15 | 15 | 15 | 13 | 15 | 8 | 15 | 15 | 15 | 15 | 15 | 15 | | | | 15 | 14 | 14 |
| *D. immigrans* | | 11 | 14 | | 15 | 12 | 15 | 15 | 15 | 13 | 14 | 13 | 12 | 15 | 13 | | 13 | 13 | 14 | 8 | 15 | 15 | 15 | 11 | 14 | 10 | 14 | 14 | 15 | 14 | 15 | 14 | 14 |
| *D. lacicola* | 14 | 12 | 11 | 13 | 15 | 12 | 15 | 12 | 15 | 12 | 11 | 14 | 15 | 13 | 14 | 14 | 12 | 13 | 13 | 11 | 10 | 14 | 14 | 15 | 15 | 14 | 14 | 15 | 13 | 14 | 15 | 13 | 14 |
| *D. melanogaster* | 15 | 15 | 15 | 13 | 15 | 14 | 15 | 13 | 13 | 13 | 15 | 15 | 15 | 15 | 15 | 15 | 15 | 15 | 15 | 14 | 14 | 15 | 14 | 15 | 15 | 15 | 15 | 15 | 15 | 15 | 13 | 14 | 15 |
| *D. montana* | 15 | 15 | 15 | 15 | 15 | 15 | 15 | 14 | 15 | 15 | 15 | 14 | 15 | 15 | 14 | 14 | 15 | 15 | 15 | 15 | 15 | 15 | 15 | 15 | 15 | 15 | 12 | 15 | 13 | 15 | 15 | 15 | 15 |
| *D. nasuta* | 15 | 13 | 15 | 12 | 14 | 15 | 15 | 14 | 12 | 12 | 14 | 14 | 12 | 14 | 12 | 12 | 14 | 15 | 12 | 14 | 14 | 14 | 11 | 13 | 14 | 11 | 14 | 15 | 14 | 14 | 9 | 14 | 15 |
| *D. nebulosa* | 14 | 8 | 13 | 15 | 10 | 8 | 10 | 8 | 7 | 11 | 14 | 9 | 13 | 12 | 12 | 13 | 11 | 11 | 11 | 11 | 12 | 15 | 9 | 8 | 12 | 13 | 14 | 14 | 10 | 10 | 11 | 11 | 10 |
| *D. paramelanica* | 15 | 13 | 15 | 13 | 14 | 14 | 13 | 12 | 13 | 11 | 15 | 14 | 12 | 14 | 14 | 11 | 11 | 12 | 13 | 12 | 14 | 12 | 13 | 13 | 15 | 14 | 15 | 14 | 12 | 14 | 14 | 13 | 14 |
| *D. persimilis* | 15 | 14 | 12 | 15 | 15 | 15 | 15 | 12 | 13 | 15 | 15 | 15 | 14 | 14 | 15 | 14 | 14 | 14 | 15 | 15 | 12 | 14 | 14 | 12 | 13 | 14 | 15 | 14 | 13 | 14 | 15 | 14 | 11 |
| *D. prosaltans* | 15 | 15 | 13 | 15 | 13 | 13 | 15 | 13 | 15 | 14 | 15 | 15 | 15 | 14 | 15 | 15 | 15 | 14 | 15 | 15 | 15 | 15 | 14 | 14 | 15 | 14 | 13 | 13 | 15 | 15 | 15 | 15 | 11 |
| *D. pseudoobscura* | 15 | 15 | 15 | 14 | 15 | 14 | 14 | 15 | 13 | 14 | 15 | 14 | 15 | 15 | 14 | 14 | 14 | 13 | 15 | 14 | 15 | 14 | 15 | 13 | 14 | 14 | 15 | 15 | 15 | 15 | 15 | 13 | 15 |
| *D. putrida* | 15 | 13 | 14 | 13 | 15 | 13 | 13 | 13 | 9 | 12 | 14 | 13 | 15 | 15 | 12 | 14 | 11 | 15 | 15 | 13 | 13 | 11 | 11 | 11 | 13 | 14 | 14 | 13 | 11 | 12 | 12 | 15 | 11 |
| *D. saltans* | 15 | 15 | 14 | 13 | 15 | 15 | 15 | 15 | 14 | 15 | 12 | 15 | 15 | 14 | 13 | 15 | 14 | 15 | 14 | 14 | 15 | 15 | 15 | 15 | 15 | 14 | 15 | 15 | 15 | 13 | 14 | 13 | 14 |
| *D. santomea* | 14 | 15 | 11 | 14 | 15 | 15 | 14 | 15 | 14 | 14 | 11 | 13 | 13 | 12 | 15 | 13 | 13 | 13 | 13 | 15 | 14 | 13 | 13 | 15 | 14 | 14 | 13 | 13 | 13 | 14 | 12 | 13 | 15 |
| *D. simulans* | 15 | 14 | 15 | 15 | 14 | 14 | 15 | 15 | 15 | 11 | 15 | 14 | 15 | 14 | 14 | 15 | 14 | 15 | 14 | 15 | 15 | 12 | 15 | 15 | 14 | 15 | 14 | 14 | 14 | 14 | 14 | 15 | 15 |
| *D. sturtevanti* | 8 | 7 | 13 | 12 | 8 | 13 | 7 | 9 | 12 | 13 | | 15 | 14 | 7 | 13 | 11 | 7 | 15 | 15 | 8 | 15 | 10 | 7 | 13 | 8 | 7 | 13 | 10 | 8 | 12 | 14 | 8 | 9 |
| *D. subobscura* | 14 | 13 | 15 | 14 | 14 | 15 | 14 | 14 | 15 | 12 | 10 | 13 | 15 | 15 | 14 | 14 | 14 | 14 | 12 | 15 | 14 | 14 | 15 | 15 | 7 | 14 | 15 | 10 | 13 | 15 | 10 | 14 | 15 |
| *D. sucinea* | 13 | 9 | 14 | 13 | 14 | 14 | 12 | 10 | 15 | 9 | 11 | 14 | 14 | 14 | 14 | 10 | 13 | 12 | 11 | 12 | 12 | 10 | 11 | 7 | 10 | 12 | 13 | 12 | 10 | 15 | 12 | 13 | 9 |
| *D. takahashii* | 15 | 15 | 15 | 11 | 15 | 15 | 13 | 15 | 15 | 15 | 13 | 15 | 15 | 15 | 15 | 15 | 13 | 15 | 15 | 15 | 15 | 9 | 15 | 14 | 13 | 15 | 15 | 15 | 15 | 14 | 12 | 15 | 15 |
| *D. teisseri* | 13 | 14 | 13 | 15 | 15 | 15 | 14 | 12 | 14 | 15 | 13 | 14 | 11 | 13 | 12 | 9 | 12 | 14 | 12 | 15 | 14 | 10 | 14 | 15 | 15 | 13 | 14 | 14 | 12 | 15 | 14 | 12 | 14 |
| *D. virilis* | 15 | 15 | 15 | 15 | 13 | 15 | 15 | 15 | 14 | 15 | 15 | 15 | 14 | 15 | 15 | 15 | 13 | 14 | 15 | 15 | 14 | 15 | 15 | 15 | 15 | 15 | 15 | | 14 | 15 | 15 | 15 | 14 |
| *D. yakuba* | 10 | 13 | 12 | 11 | 13 | 13 | 12 | 12 | 11 | 10 | 11 | 11 | 10 | 13 | 12 | 12 | 9 | 11 | 13 | 9 | 14 | 10 | 10 | 12 | 13 | 15 | 11 | 11 | 7 | 12 | 11 | 15 | 13 |
| *S. lebanonensis* | 15 | 14 | 15 | 15 | 15 | 14 | 12 | 13 | 15 | 15 | 14 | | 14 | 14 | 15 | 15 | 15 | 14 | 10 | 15 | 15 | 15 | 9 | 13 | 14 | 15 | 15 | 15 | 14 | 15 | 13 | 15 | 15 |
| *S. pattersoni* | 15 | 15 | 15 | 14 | 11 | 14 | 15 | 14 | 13 | 15 | 15 | 14 | 15 | 15 | 15 | 14 | 15 | 15 | 13 | 15 | 14 | 15 | 15 | 15 | 15 | 7 | 15 | 15 | 15 | 15 | 14 | 15 | 14 |
| *Z. davidi* | 15 | 15 | 15 | 15 | 14 | 15 | 15 | 15 | 12 | 13 | 15 | 14 | 15 | 15 | 10 | 10 | 15 | 14 | 15 | 15 | 15 | 13 | 15 | 15 | 13 | 15 | | 15 | 15 | 15 | 15 | 15 | 15 |
| *Z. tuberculatus* | 15 | 14 | 14 | 15 | 15 | 14 | 15 | 14 | 15 | 15 | 15 | 15 | 15 | 15 | 15 | 14 | 15 | 14 | 14 | 15 | 13 | 14 | 14 | 14 | 12 | 13 | 15 | 15 | 15 | 15 | 13 | 14 | 14 |

**Supplementary Table 2: Biological replicates (vials) and flies per vial by species and virus isolate.** Black boxes indicate replicates that were excluded from analysis, all of which were removed due to melt-curve contaminants.

| Isolate | NCBI Accession |
|---|---|
| CrPV-BEE | PQ246907 |
| CrPV-GRA | PQ246908 |
| CrPV-KKH | PQ246909 |
| CrPV-KTA | PQ246910 |
| CrPV-NEU | PQ246911 |
| CrPV-OG | PQ246912 |
| CrPV-VIC | PQ246913 |
| CrPV-WHP | PQ246914 |
| DCV-C | MK645242.1 |
| DCV-CYG | MK645238.1 |
| DCV-EB | MK645239.1 |
| DCV-G | MK645241.1 |
| DCV-M | MK645243.1 |
| DCV-O | MK645244.1 |
| DCV-T | MK645245.1 |
| DCV-Z | MK645240.1 |

**Supplementary Table 3: DCV and CrPV isolate NCBI Genome Accessions**

| Virus | Forward | Reverse |
|---|---|---|
| CrPV | GGAGAACCGATTTCGTATGA | GTTGGTGGAATGTCTTCTCT |
| DCV | GACACTGCCTTTGATTAG | CCCTCTGGGAACTAAATG |
| FHV | TTATTATGTCACCGAGCCTG | CTTCGGGTTAAAGGTGTGTA |
| DAV | TATCTTACCAAAGGCAACCC | CAAACTCAATCACCCATTCG |
| IIV6 | GAACACAACAAACCGTTTCC | GGTGCAGATGGTGTAACAAT |
| DmelNv | TGGTTTGTATGCGTGGGTGA | GCTCGAGACATTCTGTCGGT |
| BFV | ATGTTGACTCGACGTACTAC | GCGTATTTCAAAGCATGACA |

**Supplementary Table 4: Viral q-PCR primers**

| Direction | Name | Sequence |
|---|---|---|
| Forward | F-a | TGCCAAGTTGTCGCACAAATGG |
| | F-b | TGCTAAGTTGTCGCACAAATGG |
| | F-c | TGCCAAGCTGTCGCACAAATGG |
| | F-d | TGCTAAGCTGTCGCACAAATGG |
| | F-e | TGCGAAGTTGTCGCACAAATGG |
| | F-f | TGCGAAGCTGTCGCACAAATGG |
| Reverse (cDNA) | R-a | TGCGCTTGTTGGAACCGTAAC |
| | R-b | TGCGCTTGTTGGATCCGTAAC |
| | R-c | TGCGCTTGTTGGAACCATAAC |
| | R-d | TGCGCTTGTTGGAGCCGTAAC |
| | R-e | TGCGCTTGTTAGAACCGTAAC |
| | R-f | TACGCTTGTTGGAACCGTAAC |
| | R-g | TGCGCTTGTTGGAACCGTAGC |
| | R-h | TGCGCTTGTTCGATCCGTAAC |
| | R-i | TGCGCTTGTTGGAGCCATAAC |
| | R-j | TGCGCTTGTTTGATCCGTAAC |
| | R-k | TGCGCTTGTTTGAACCATAAC |
| | R-l | TACGCTTGTTGGAACCATAAC |
| | R-m | TACGCTTGTTGGAGCCGTAAC |
| | R-n | TGCGCTGGTTGGAACCATAAC |
| | R-o | TGAGCTTGTTCGATCCGTAAC |
| | R-p | TACGCTTGTTGGAGCCATAAC |
| | R-q | TGAGCTTGTTTGATCCGTAAC |
| | R-r | TAAGCTTGTTGGATCCGTAGC |
| | R-s | TCAGCTTGTTGGATCCATAGC |
| Reverse (gDNA) | R-gDNA | GGYTTRCGCCATTTGTGC |

**Supplementary Table 5: RPL32 q-PCR primers**

| Species | Forward | Reverse (cDNA) | Reverse (gDNA) |
|---|---|---|---|
| *D. affinis* | F-a | R-i | R-gDNA |
| *D. americana* | F-c | R-a | R-gDNA |
| *D. ananassae* | F-f | R-a | R-gDNA |
| *D. arizonae* | F-a | R-a | R-gDNA |
| *D. baimaii* | F-a | R-r | R-gDNA |
| *D. buzzati* | F-a | R-e | R-gDNA |
| *D. erecta* | F-d | R-h | R-gDNA |
| *D. euronotus* | F-a | R-g | R-gDNA |
| *D. flavomontana* | F-c | R-a | R-gDNA |
| *D. hydei* | F-a | R-a | R-gDNA |
| *D. immigrans* | F-b | R-p | R-gDNA |
| *D. lacicola* | F-c | R-a | R-gDNA |
| *D. melanogaster* | F-d | R-h | R-gDNA |
| *D. montana* | F-c | R-a | R-gDNA |
| *D. nasuta* | F-b | R-f | R-gDNA |
| *D. nebulosa* | F-b | R-c | R-gDNA |
| *D. paramelanica* | F-a | R-g | R-gDNA |
| *D. persimilis* | F-a | R-b | R-gDNA |
| *D. prosaltans* | F-a | R-n | R-gDNA |
| *D. pseudoobscura* | F-a | R-m | R-gDNA |
| *D. putridia* | F-d | R-q | R-gDNA |
| *D. saltans* | F-a | R-n | R-gDNA |
| *D. santomea* | F-a | R-n | R-gDNA |
| *D. simulans* | F-d | R-h | R-gDNA |
| *D. sturtevanti* | F-a | R-l | R-gDNA |
| *D. subobscura* | F-a | R-i | R-gDNA |
| *D. sucinea* | F-b | R-k | R-gDNA |
| *D. takahashii* | F-d | R-o | R-gDNA |
| *D. teissieri* | F-d | R-h | R-gDNA |
| *D. virilis* | F-c | R-a | R-gDNA |
| D. yakuba | F-d | R-h | R-gDNA |
| S. lebanonensis | F-d | R-h | R-gDNA |
| S. pattersoni | F-a | R-m | R-gDNA |
| Z. davidi | F-a | R-c | R-gDNA |
| Z. tuberculatus | F-a | R-c | R-gDNA |

**Supplementary Table 6: RPL32 Primer Combinations**

| Cycle step | Temp. | Time/rate | Cycle No. |
|---|---|---|---|
| Initial denaturation | 95˚C | 2 min | 1 |
| Denaturation | 95˚C | 5 sec | } 40 |
| Annealing/Extension | 60˚C | 15 sec | |
| Melt curve | 60˚C - 95˚C | 0.1˚C/s | 1 |

**Supplementary Table 7: q(RT)-PCR Cycle Conditions**

| Virus | Repeatability | Phylogenetic Heritability |
|---|---|---|
| CrPV-OG | **0.84 (0.74, 0.93)** | 0.47 (0.00, 0.97) |
| CrPV-GRA | **0.69 (0.51, 0.86)** | **0.73 (0.19, 1.00)** |
| CrPV-VIC | **0.82 (0.71, 0.92)** | **0.75 (0.13, 1.00)** |
| DCV-C | **0.90 (0.82, 0.96)** | **0.72 (0.23, 1.00)** |
| DCV-EB | **0.96 (0.93, 0.98)** | **0.86 (0.52, 1.00)** |
| DCV-M | **0.87 (0.79, 0.94)** | **0.87 (0.59, 1.00)** |
| FHV | **0.22 (0.04, 0.42)** | **0.85 (0.39, 1.00)** |
| DAV | **0.62 (0.4, 0.82)** | **0.79 (0.35, 1.00)** |
| IIV6 | **0.77 (0.64, 0.88)** | **0.95 (0.78, 1.00)** |
| DmelNV | **0.57 (0.38, 0.75)** | **0.94 (0.77, 1.00)** |
| BFV | 0.17 (0, 0.41) | 0.54 (0.01, 1.00) |

**Supplementary Table 8: Estimates of repeatability and phylogenetic heritability in viral load for each virus.** All models were fitted on $log_{10}$ transformed fold-changes in viral load. Values for phylogenetic heritability are taken from a model containing a non-phylogenetic species-specific random effect, while values for repeatability are taken from a model with this effect removed. Estimates that are credibly non-zero are shown in bold.

| Group | Correlation Coefficient (R) |
|---|---|
| Within Species | 0.93 (0.82, 0.99) |
| Within Family | 0.73 (0.49, 0.93) |
| Across Family | 0.33 (-0.18, 0.79) |

**Supplementary Table 9: Estimates of the strength of correlations between viruses at different evolutionary scales.** Posterior distributions of the correlation coefficients for each virus pair were combined to create overall distributions for correlations within virus species, within virus family, and across virus family.

| Group | Correlation Coefficient (R) |
|---|---|
| Within Species | 0.93 (0.82, 0.99) |
| Within Family | 0.73 (0.49, 0.93) |
| Across Family | 0.27 (-0.11, 0.63) |

**Supplementary Table 10: Estimates of the strength of correlations between viruses at different evolutionary scales with BFV included.**

### Supplementary Figures

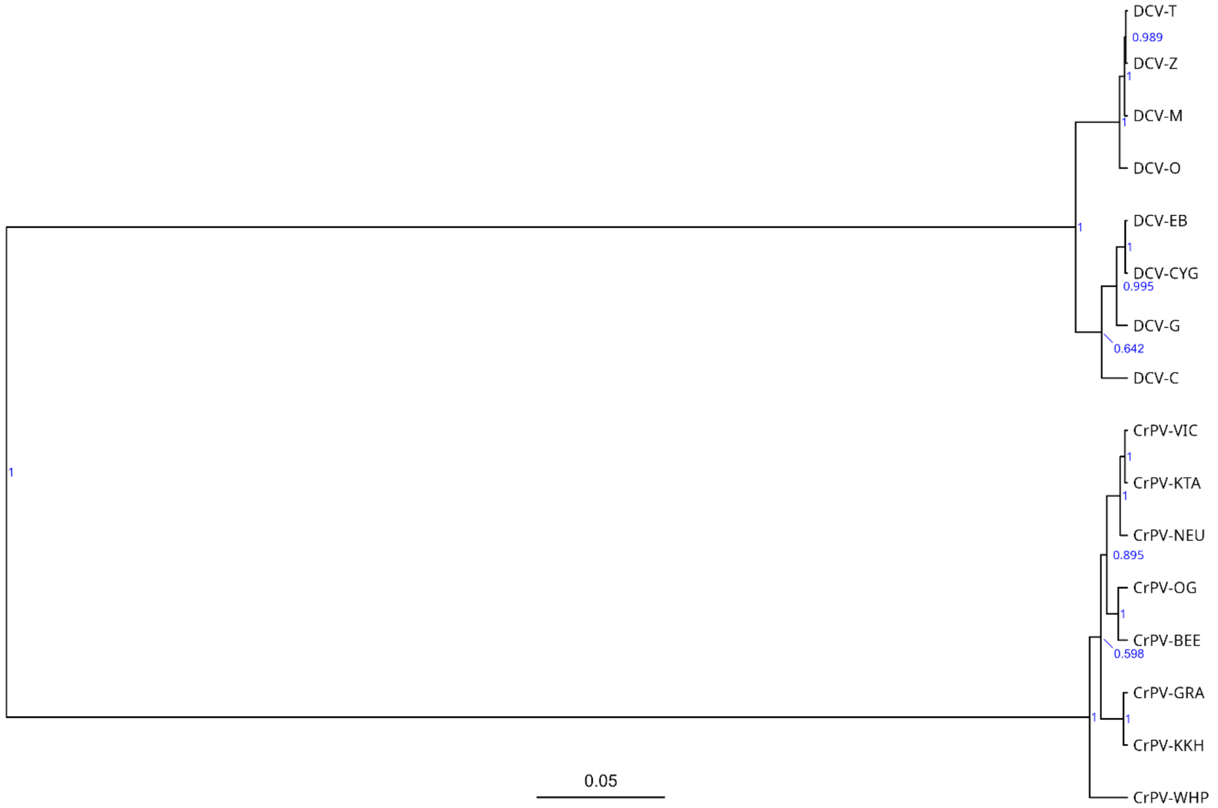


**Supplementary Figure 1: Phylogeny of DCV and CrPV isolates.** Evolutionary relationships are presented as a midpoint rooted, maximum clade credibility tree. Node labels (blue) are the posterior probabilities of each clade, and the scale bar represents nucleotide substitutions per site.

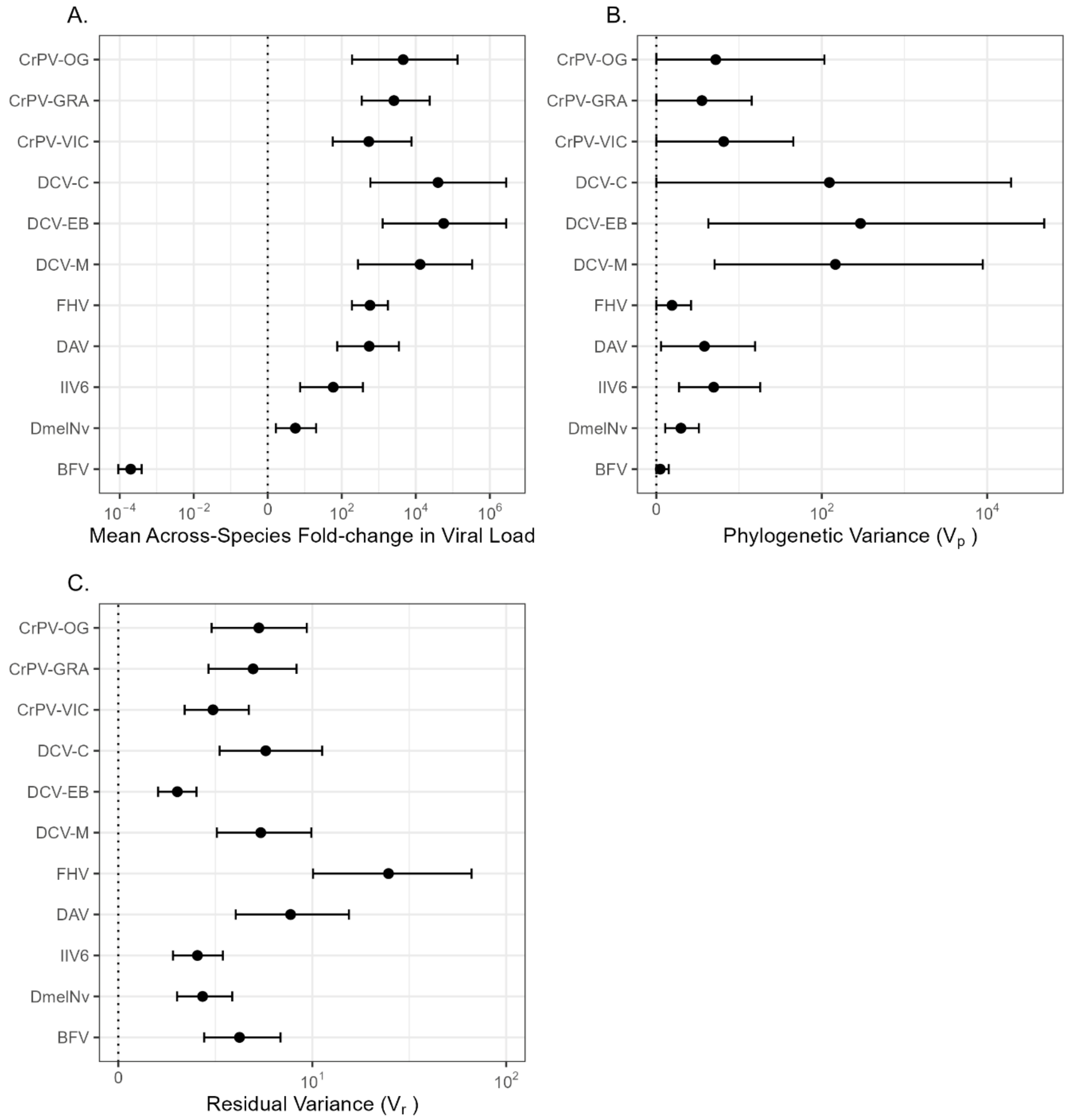


**Supplementary Figure 2: Estimates of the mean across species viral loads and variances for each virus.** Values for the mean across species fold-change in viral load (A), phylogenetic variance (B), and residual variance (C) were taken from univariate models with the non-phylogenetic species-specific random effect removed.
